# OS2CR-Diff: A Self-Refining Diffusion Framework for CD8 Imputation from One-Step Inference to Conditional Representation

**DOI:** 10.1101/2025.11.09.687488

**Authors:** Xingnan Li, Priyanka Rana, Tuba N Gide, Nurudeen A Adegoke, Yizhe Mao, James S Wilmott, Sidong Liu

## Abstract

Stain imputation in multiplex immunofluorescence (mIF) imaging addresses the challenge of missing or damaged biomarker channels by reconstructing target biomarker images from a limited set of available stains. This approach offers a faster and more efficient alternative to full-panel staining, enabling detailed analysis of the tumour microenvironment. Existing One-Step Inference Models (OSIMs), primarily based on generative adversarial networks (GAN) or autoencoders, often generate suboptimal images with significant artifacts or reduced signal intensity. These limitations impair visual interpretability and reliability of the downstream immunotherapy response assessment. The challenge is further amplified when imputing cytoplasmic biomarkers such as CD8 from commonly used stains such as DAPI, due to the limited spatial correlation and the inherently complex structure of cytoplasmic signals. To address these limitations, we propose a self-refining diffusion model, OS2CR-Diff, which utilises the results from OSIMs as additional conditional representations. Unlike prior studies that rely on a single or limited conditional inputs, OS2CR-Diff incorporates three conditional inputs: the OSIM-imputed target biomarker image, OSIM-imputed complementary biomarker images, and non-antibody-stained images. Furthermore, we propose a feature fusion module that employs a cross-gated attention mechanism to effectively integrate these inputs, enabling context-aware feature refinement and improving the quality and reliability of imputed biomarker images. We evaluated OS2CR-Diff for CD8 biomarker imputation on mIF images of melanoma tissues. Our method out-performed state-of-the-art methods, achieving a 73.4% increase in the Structural Similarity Index Measure (SSIM), a 28.9% gain in the Peak Signal-to-Noise Ratio (PSNR), a 61.2% improvement in Mean Absolute Error (MAE), and significantly lower false positive rates compared to OSIM.

## I. Introduction

**M**Ultiplex immunofluorescence (mIF) imaging generates rich and multidimensional data that capture the simultaneous expression patterns of multiple biomarkers. This enables a detailed analysis of the tumour microenvironment, supports spatial profiling of biomarkers, and advances research in immuno-oncology [1]. Among these biomarkers, nuclear biomarkers, such as Ki67 and FOXP3, indicate cell proliferation or cell cycle status [2], while cytoplasmic markers reflect functional processes such as cytokine production and intracellular signalling, providing a more direct link to the immune and therapeutic response [3]. Specifically, CD8 is one of the most robust and measurable cytoplasmic biomarkers [4], as it is essential for the reliable assessment of immune cell infiltration, a key factor in evaluating the patient’s response to immunotherapy, particularly in melanoma [5]. mIF enhances this assessment by enabling the detection of spatial interactions between cells, which are closely associated with clinical outcomes [6], [7].

mIF imaging involves staining multiple biomarkers within a single tissue sample and faces significant technical challenges, including an increased risk of procedural failure, staining variability, and channel dropout [8]. These issues often result in missing or low-quality biomarker channels. Machine learning models trained on inaccurate or incomplete images suffer from degraded performance, potentially causing significant errors in clinical assessment. This emphasises the critical need for stain imputation methods that can reconstruct missing or corrupted biomarker signals, thereby ensuring high-quality, accurate imaging data for reliable model training and robust immune cell analysis.

Accordingly, recent advances in Generative Adversarial Networks (GANs) [9] and Vision Transformers (ViTs) [10] have enabled the development of One-Step Inference Models (OSIMs) as practical alternatives to reconstruct incomplete biomarker images through stain imputation. Specifically, models such as Stain Imputation in Multiplex Immunofluorescence Imaging (SIMIF) [11], Multimodal-Attention-based virtual mIF Staining (MAS) [12] generate biomarker images, such as CD8 and PD-L1, from readily available non-antibody-stained biomarker images (DAPI).

However, these models struggle to produce accurate and artifact-free imputation results, particularly for cytoplasmic biomarker images due to their inherently complex structure. The imputed images often exhibit blurred signals, inaccurate intensity predictions, and extensive artifacts. This is due to fundamental differences in staining targets: DAPI stains the inner-nuclear region [13], while cytoplasmic biomarkers highlight the outer-nuclear region [14]. This mismatch in staining patterns presents a substantial challenge for OSIMs, limiting their clinical and research utility.

To enhance the visual quality of OSIM-generated images, Liu et al. [15] proposed a Schrödinger Bridge image-to-image (I2SB) denoising diffusion probabilistic model (DDPM) [16]. This model directly learns the nonlinear diffusion processes from degraded images to high-resolution outputs and has shown improved performance in natural image translation tasks. Additionally, Zhou et al. [17] proposed Cascade Multi-path Shortcut Diffusion Model (CMDM) to improve the translation performance of medical images by using a diffusion model to improve the GAN-generated image. However, both methods refine the OSIM generated images that are often suboptimal in quality. Consequently, any inaccuracies in these inputs may be propagated or even amplified by the diffusion model, potentially leading to degraded image quality and reduced reliability in downstream analysis and clinical decision-making. Furthermore, current approaches have not explored incorporating additional conditional inputs, such as OSIM-imputed biomarker images that are complementary to the target biomarker to enrich contextual information and enhance performance.

Accordingly, to support clinical interpretation and downstream analysis, we propose a self-refining diffusion model that incorporates the outputs from OSIM as additional conditional representations (OS2CR-Diff), aiming to enhance biomarker images initially imputed by OSIM. It builds on the outputs generated by MAS [12], an OSIM capable of simultaneously imputing multiple biomarker images from non-antibody stains (DAPI and autofluorescence (AF)). OS2CR-Diff includes a novel conditional feature fusion block that integrates multiple conditional inputs and employs a U-Net-based DDPM backbone [18]. Furthermore, to accelerate convergence and prevent brightness shifts, we employ velocity prediction (*v*-prediction) [19] instead of noise parameterisation (*ε*-prediction). Given the critical role of CD8 in the immunotherapy response, we evaluated our proposed method using CD8 biomarker images.

In summary, our main contributions are as follows:

- Multi-conditional input design for context-aware generation: Unlike existing methods, our framework integrates three conditional inputs: OSIM-imputed target biomarker, OSIM-imputed complementary biomarker images and non-antibody-stained images. This multi-source conditioning enables more context-aware generation and refinement of target biomarker images.
- Conditional feature fusion block: We design a fusion block that integrates OSIM-imputed target and complementary biomarkers with non-antibody stains using a cross-gated attention mechanism [20] for effective feature fusion.
- Visual quality improvement: The proposed method improves the quality of cytoplasmic biomarker images (e.g., CD8) through semantic refinement, artifact reduction, and structural reconstruction. These enhancements are particularly notable given the inherent difficulty of imputing cytoplasmic biomarkers, and are especially valuable for supporting accurate downstream analysis and clinical interpretation.
- Robust CD8^+^ T cells counting performance: Evaluation of CD8^+^T cells detection and segmentation demonstrates that imputed images significantly reduce false positive rates, thereby providing more accurate and clinically reliable images for assessing immunotherapy response.

## II. Related work

### A. Stain imputation methods for mIF images

Stain imputation has emerged as a pivotal advancement in mIF imaging, offering the potential to streamline workflows by eliminating complex staining protocols and minimising staining failures. The Marker Imputation Model for Multiplex Images (MAXIM) [21] and SIMIF [11] represent some of the initial efforts in stain imputation, utilising GAN-based architectures to generate a target biomarker image from other available markers. However, MAXIM requires retraining for specific input-output pairing, which limits its adaptability across diverse biomarker configurations. SIMIF addresses this limitation by supporting flexible input combinations while incorporating techniques that enhance training stability. Recently, MAS [12] has employed attention-based mechanisms to enable the simultaneous imputation of multiple biomarkers, offering improved flexibility and scalability. Although MAS employs an attention-based architecture for efficient multimarker prediction, computational limitations restrict it to a limited set of input biomarkers (DAPI and AF), thereby constraining its applicability.

Importantly, these models remain limited in their ability to accurately impute biomarkers, particularly cytoplasmic biomarkers such as CD8, from only non-antibody-stained inputs. The imputation of cytoplasmic biomarkers is challenging due to insufficient conditional inputs and the inherent morphological complexity of cytoplasmic structures. This limitation poses significant challenges for reliable visual interpretation in real-time clinical applications, where diagnostic precision is crucial [22].

### B. Self-refining diffusion models

Self-refining diffusion models leverage initial structural priors, such as OSIM-generated images, to guide and iteratively enhance the quality of generated outputs, surpassing the original OSIM results. Consequently, diffusion-based frameworks have been developed to refine degraded inputs and produce high-resolution, detail-preserving images. For example, Palette [23] introduced an image-conditioned diffusion architecture for guided image translation, while I2SB [15] used suboptimal OSIM-generated images as direct input for refinement. CMDM [17] further advanced the field by leveraging OSIM outputs as initialisation for medical image enhancement, effectively reducing refinement errors.

However, both CMDM and I2SB share a fundamental limitation in their reliance on the quality of the initial outputs (OSIM-generated images). Inaccurate or artifact-laden OSIM outputs can be inadvertently amplified by the diffusion process rather than corrected. Palette employs simplistic strategies, such as direct addition or concatenation, to integrate target biomarker channels with non-antibody inputs, which are inadequate for effectively handling multiple signals. The combination of limited conditional inputs and this basic fusion approach constrains Palette’s ability to integrate complex, heterogeneous inputs, which is an essential requirement in medical imaging scenarios, particularly for the mIF stain imputation task.

Therefore, the accurate reconstruction of cytoplasmic biomarkers from limited, yet effectively integrated inputs remains essential for advancing the clinical applicability of stain imputation in mIF workflows

## III. Materials and methods

### A. Dataset

This study uses a private dataset of 83 whole-slide (WS) mIF images of melanoma patients provided by Melanoma Institute Australia (MIA) [24], [25]. Each image comprises seven channels, representing six biomarkers (CD68, SOX10, CD16, CD8, PD-L1, DAPI) and one autofluorescence (AF) channel. The images have a resolution of up to 38,000 *×* 37,000 pixels, scanned at a pixel size of 0.5 *µ*m/pixel. We used 42 samples for model training and another 41 samples for testing, which were collected using different staining protocols.

### B. Data preprocessing

The images were normalised to ensure consistent intensity levels across channels. Segmentation of the tissue region is performed using an adaptive threshold based on Otsu’s method [26]. Given the uniform expression levels of key biomarkers, we used PD-L1, CD8 and DAPI to generate masks, with the DAPI mask providing contextual information to guide the extraction of region-of-interest boundary coordinates.

The segmented tissue regions were further subdivided into 224 *×* 224 pixel image patches/instances, with each WS mIF image contributing up to 1, 000 instances, yielding a final dataset of 79, 192 image instances. To prevent information leakage, all patches from the same WS mIF image and patient were assigned to the same fold.

### C. One-step-to-conditional refinement diffusion model (OS2CR-Diff)

OS2CR-Diff enhances OSIM-generated images, which are currently constrained by substantial structural and morphological discrepancies between the target biomarker and limited conditional inputs (Fig. 1). The proposed model enables precise visual inspection and significantly reduces the false-positive cell detections, thereby minimising the risk of erroneous clinical decisions and unnecessary treatments. Specifically, OS2CR-Diff, a conditional DDPM, facilitates more context-aware image generation by incorporating multiple conditional inputs. To guide accurate stain imputation, the fusion block integrates these conditional inputs using a cross-gated attention mechanism that captures interdependencies between target and auxiliary inputs, facilitating effective feature refinement.

**Fig. 1:**
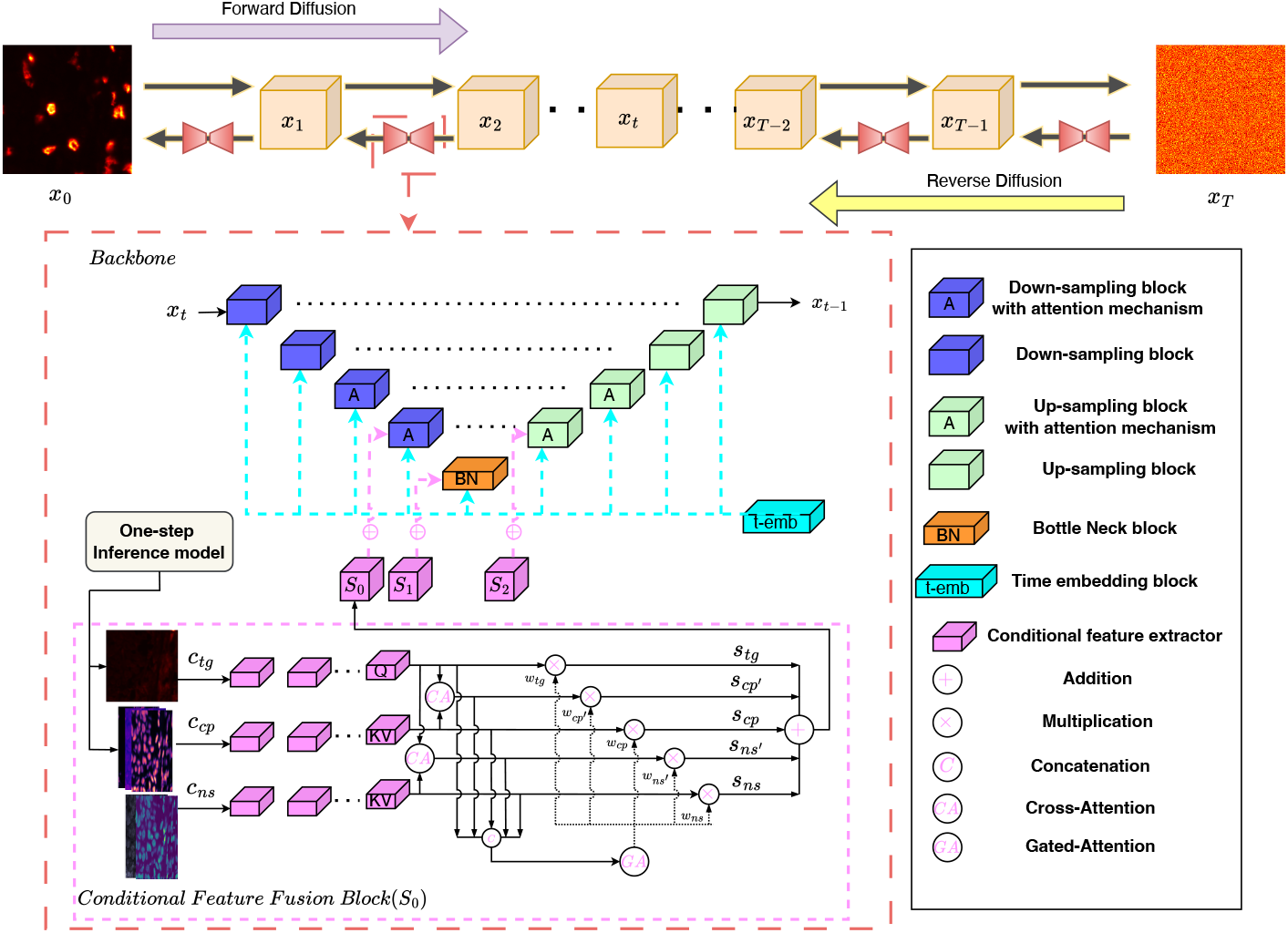
A schematic view of the proposed OS2CR-Diff for CD8 imputation in mIF imaging.

During training, random noise is progressively added to the original target biomarker images following a predefined linear diffusion schedule over T timesteps. The original biomarker images serve as ground-truth references for the model’s denoising task with a U-Net model, containing four down-sampling blocks, four upsampling blocks and one bottleneck block. Additionally, the proposed OS2CR-Diff employs *v*-prediction instead of conventional noise prediction to enhance robustness against imaging artifacts commonly encountered in clinical samples, including variations in brightness, contrast, and colour.

During inference, the OS2CR-Diff model refines the previous OSIMs-imputed target biomarker images to accurate, biologically representative images. Specifically, the inference pipeline iteratively reverses the diffusion process over T timesteps, progressively reducing noise to yield refined images. The proposed conditional feature fusion block provides enriched conditional information during this reverse process, supporting accurate image generation. Consequently, the resulting high-quality images enhance visual interpretability and contribute to informed treatment decisions.

### D. Velocity prediction parameterisation

Standard DDPMs typically employ either *ε* or *x*_0_-prediction, but both methods suffer from a mismatch between training and inference conditions, often resulting in noticeable artifacts such as mid-tone bias in grayscale images or unnatural colour shifts in RGB images [27].

To improve training stability and sample quality, recent DDPMs employ *v*-prediction, a technique that reframes the denoising objective to predict the velocity vector (a linear interpolation of the clean image and the noise), instead of directly predicting the noise or the clean image. Inspired by Stable Diffusion [28], we adopt this approach to reduce leakage shortcuts from residual low-frequency structures and to address the signal-to-noise ratio (SNR) imbalance. This results in accelerated convergence and more accurate pixel-level reconstruction, making it particularly beneficial for medical imaging applications.

Specifically, the forward diffusion process gradually corrupts a clean image *x*_0_ at time step t as:

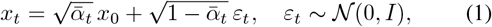

where *x*_*t*_ denotes the noisy intermediate image obtained at diffusion step t and 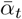 denotes the cumulative product of noise-scheduling coefficients. The velocity vector *v*_*t*_ is explicitly defined as:

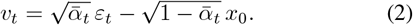

During training, our proposed model predicts the velocity 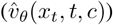, which incorporates conditional information *c* and is optimised with all trainable parameters (*θ*) using a mean squared error (MSE) loss function:

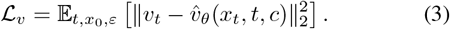

During inference, high-fidelity reconstruction is achieved by reversing the diffusion process: at each timestep, the predicted velocity 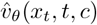 is algebraically transformed into the necessary intermediate quantities to compute the posterior mean and variance.

### E. Conditional feature fusion block

To enrich spatial context and guide more accurate stain imputation, we effectively incorporate multiple sources of conditional information into our model, including *c*_*tg*_, *c*_*cp*_, and *c*_*ns*_. Here, *c*_*tg*_ represents OSIM-generated target images, *c*_*cp*_ represents OSIM-generated complementary biomarker images, and *c*_*ns*_ refers to non-antibody-stained images. Accordingly, we introduce a conditional feature fusion block designed to independently extract feature representations from each conditional input. Within this block, the cross-attention mechanisms are employed to uncover latent interdependencies between the target and auxiliary conditions to refine features. A subsequent gated attention module adaptively weights the conditional inputs, enabling context-aware fusion within the U-Net latent space. These enriched features enhance the model’s ability to emphasise relevant information during inference, ultimately producing more informative representations for biomarker imputation.

We strategically merge conditional features at three key locations within the U-Net: the last downsampling block, the first upsampling block, the bottleneck, as shown in Fig. 1. Each fusion block incorporates three independent feature extractors to process the conditional inputs (*c*_*tg*_, *c*_*cp*_, and *c*_*ns*_). For each conditional input, we use several convolutional layers to map features to the appropriate dimension, for example 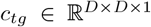 is transformed to 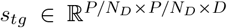, where *P* represents the original image size, *N*_*D*_ represents the number of downsampling operations, and *D* represents the corresponding dimension at the U-Net block. The same feature extraction process is applied to *c*_*cp*_ and *c*_*ns*_ to obtain *s*_*cp*_ and *s*_*ns*_, respectively. This procedure is repeated for all three fusion locations within the network.

Furthermore, we implement a cross-attention mechanism using *s*_*tg*_ as the query, with each of the other two features serving as keys and values to generate refined representations *s*_*cp*_′ and *s*_*ns*_′ . The cross-attention between *s*_*tg*_ and *s*_*cp*_ enables the target biomarker to capture spatial correlation information from complementary biomarkers. This improves the ability of the model to estimate the true localisation of the target biomarker during the denoising process. Meanwhile, the crossattention between *s*_*tg*_ and *s*_*ns*_ aims to refine latent correlations and leverage structural cues from the reference channels. This provides a coarse yet essential spatial prior for subsequent refinement steps.

After obtaining the two refined feature representations through the cross-attention mechanism (*s*_*cp*_′ and *s*_*ns*_′ ), we incorporate them along with the other three conditional features using a gated attention mechanism. This mechanism enables the model to adaptively assign attention weights (*w*_*tg*_, *w*_*cp*_′, *w*_*cp*_, *w*_*ns*_′, and *w*_*ns*_) to each of the five inputs (*s*_*tg*_, *s*_*cp*_′, *s*_*cp*_, *s*_*ns*_′, and *s*_*ns*_), based on their relevance to the target task. By dynamically controlling the flow of information from each conditional source, the gated attention mechanism improves the contextual richness and specificity of the final conditional representation, which enhances the performance of the denoising diffusion process. The final fused feature (*S*) is obtained through weighted addition, providing enhanced conditional representations for the denoising procedure.

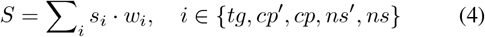

where *S* represents features that need to be fused into the last downsampling block, the first upsampling block, and the bottleneck of the U-Net, respectively, as shown in Fig. 1. In Figure 1, the variable *S* is renamed depending on the block: *S*_0_ for the downsampling block, *S*_2_ for the upsampling block, and *S*_1_ for the bottleneck block.

### F. Implementation

All diffusion-based models were trained for over 500,000 steps using a linear beta schedule for the forward diffusion process, where the noise schedule (*β*_*t*_) increases linearly from 1 *×* 10^−4^ to 2 *×* 10^−2^ over 1000 timesteps, following the original DDPM formulation. Additionally, all OSIMs, including MAXIM, SIMIF, and MAS, followed the training schedules defined in their respective studies. All models were trained using a learning rate of 5 *×* 10^−5^, the Adam optimiser, and a batch size of 16 on Tesla V100 GPUs.

## IV. Results

We evaluated the performance of OS2CR-Diff on the dataset containing 41 mIF samples using Mean Absolute Error (MAE) [29], Peak Signal-to-Noise Ratio (PSNR) [30], and Structural Similarity Index Measure (SSIM) [30]. We note that the metrics are computed at the WS level instead of the instance level to mitigate biases arising from the heterogeneous pixel distribution within the WS. Due to the clinical relevance and challenges in the imputation of CD8 biomarkers, we evaluated our method on CD8-stained images. The three conditional inputs used are: CD8 (target biomarker), CD16, CD68, and SOX10 (complementary biomarkers), and DAPI and AF (non-antibody biomarkers). Ablation studies were performed to assess the contributions of different conditional input configurations and the proposed feature fusion module within the proposed framework.

To further evaluate the utility of the imputed CD8 images for downstream tasks, we performed cell segmentation using Cellpose-SAM [31] and measured improvements in positive cell detection over OSIM-imputed images using the Dice coefficient [32].

### A. Comparison with state-of-the-art methods

We evaluated the performance of OS2CR-Diff against current state-of-the-art (SOTA) one-step inference stain imputation methods, including MAXIM [21], SIMIF [11], and MAS [12], as well as recent self-refining diffusion models such as Palette [23] and CMDM [17] for the imputation of CD8 image samples (Table I). The results are analysed both quantitatively, qualitatively (visual inspection).

**TABLE I:** Performance comparison for CD8 imputation from non-antibody-stained biomarkers.

| Method | Conditional Inputs |  |  | Imputed CD8 |  |  |
| --- | --- | --- | --- | --- | --- | --- |
|  | DAPI & AF | One-step-Imputed CD8 | One-step-Imputed CD16, CD68, Sox10 | MAE ↓ | SSIM ↑ | PSNR ↑ |
| MAXIM [21] | ✓ | × | × | 57.97 | 0.007 | 11.77 |
| SIMIF [11] | ✓ | × | × | 26.76 | 0.312 | 18.68 |
| MAS [12] | ✓ | × | × | 25.22 | 0.334 | 17.68 |
| Palette [23] | ✓ | ✓ | × | 12.11 | 0.512 | 21.13 |
| CMDM [17] | ✓ | ✓ | × | 20.11 | 0.432 | 20.01 |
| **OS2CR-Diff (our method)** | ✓ | ✓ | ✓ | **9.78** | **0.579** | **22.79** |

Among the three OSIMs evaluated, MAXIM shows the lowest performance across all metrics, due to its inflexible model configuration, which prevents it from effectively capturing meaningful biomarker patterns from only non-antibody-stained inputs. In contrast, SIMIF (which accommodates flexible inputs) and MAS (which employs an attention based stain imputation method) achieve comparable results, as both utilise identical conditional inputs (DAPI and AF) during the inference phase. In particular, compared to the best performing and flexible OSIM (MAS), OS2CR-Diff improved SSIM by 73.4%, PSNR by 28.9%, and reduced 61.2% by MAE. Furthermore, OS2CR-Diff outperformed current state-of-the-art self-refining models (Palette and CMDM), delivering a 13.1% increase in SSIM, a 19.2% reduction in MAE, and a 7.9% improvement in PSNR.

Fig. 2 highlights three distinct improvements offered by our proposed method over existing approaches: enhanced semantic representation, effective artifact removal, and improved structural reconstruction. We note that the GAN-based models, such as MAXIM and SIMIF, suffer from substantial signal loss and visual artifacts (third and fourth rows from the top). In contrast, the corresponding results from the transformer-based model MAS, alleviate some of these issues; however, noticeable artifacts remain, especially when the imputed images are viewed at higher magnification. These distortions may mislead clinical interpretation, especially in the context of immunotherapy response assessment.

**Fig. 2:**
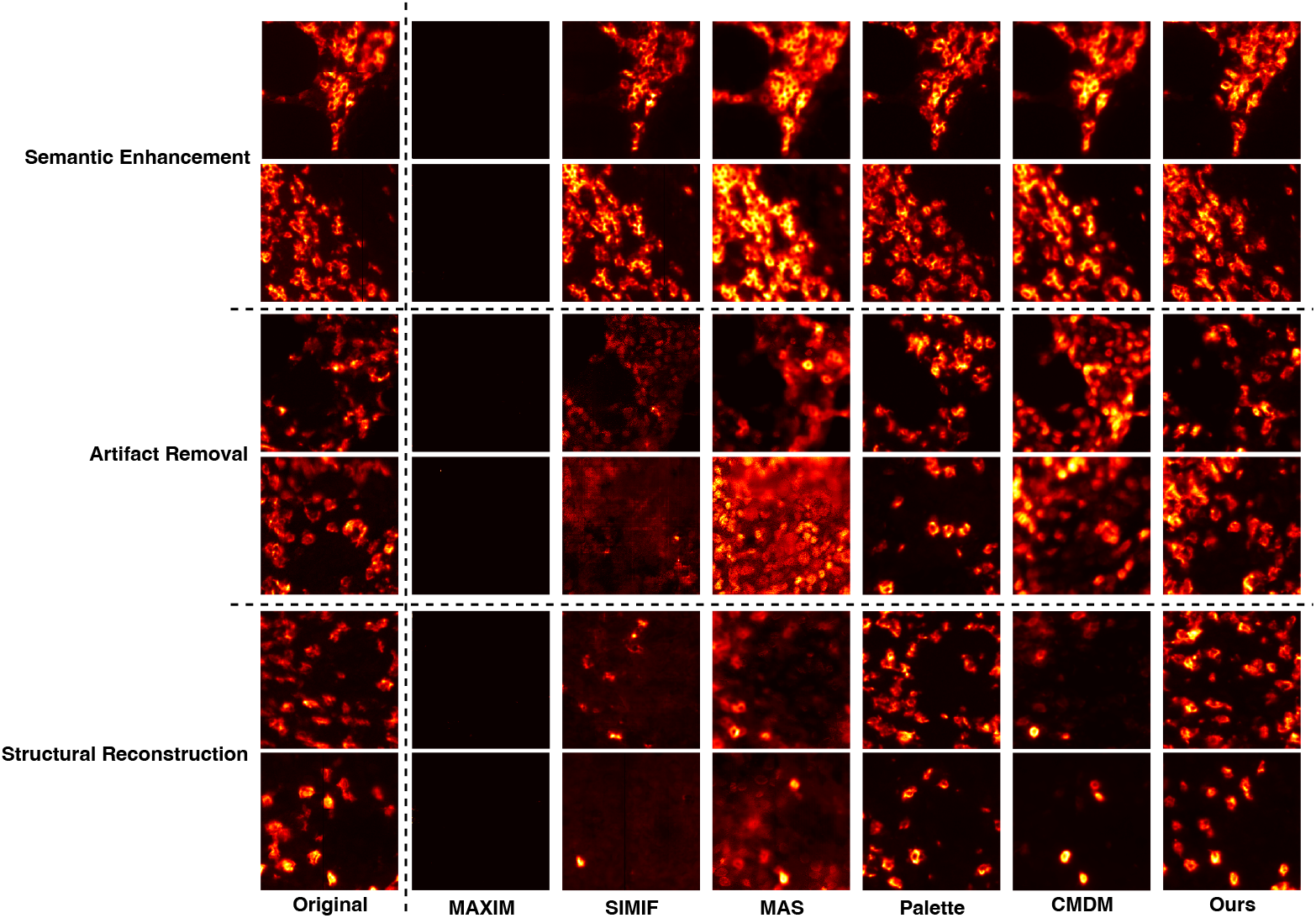
Visualisation of original and imputed images for CD8 in melanoma. From top to bottom, the first two rows illustrate enhanced semantic information in the refined CD8 images; the third and fourth rows highlight the reduction of generated artifacts; and the last two rows demonstrate improved structural details in the imputed images

A common limitation of OSIMs, is insufficient conditional information and reliance on pixel intensity averaging to impute corresponding regions, which can diminish the diversity of biomarker signals. Moreover, their exclusive reliance on DAPI and AF inputs may encourage the model to reproduce patterns from these reference stains, rather than learning meaningful biomarker-specific representations.

Furthermore, although the CMDM-imputed images (Fig. 2) exhibit pixel-wise visual improvements, they yield less accurate results than OS2CR-Diff due to a mismatch between the subcellular localisation of the target biomarkers and the conditioning input. Similarly, Palette-imputed images often show denoising artifacts, which make it difficult to accurately locate cells and can lead to false identification of CD8^+^T cells. The results (Fig.2, Table I) demonstrate that our model significantly enhances the structural semantic details in the refined CD8 and effectively reduces the artifacts generated by previous methods.

The three key improvements achieved by OS2CR-Diff (semantic enhancement, artifact removal, and structural reconstruction) are attributed to the proposed method’s effective integration of three conditional inputs. Non-antibody stains capture spatially aligned cellular structures, with DAPI providing strong nuclear localisation as a structural guide and AF contributing additional contextual information from autofluorescence signals. Complementary biomarkers provide high-level semantic context, enabling the model to perform informed imputation even in regions with missing or degraded target signals. The OSIM-imputed CD8 target biomarker serves as a prior, guiding the model to focus on relevant regions. To effectively integrate these diverse features, a cross-gated attention mechanism is used, which selectively emphasises informative signals while suppressing irrelevant noise.

### B. Ablation studies

To evaluate the contribution of individual components in OS2CR-Diff, we conducted ablation studies to understand how different combinations of conditional inputs and attention mechanisms affect CD8 image imputation performance (Table II). Specifically, three configurations were evaluated: (i) a base configuration, which used only DAPI and AF channels, along with the MAS imputed CD8 image as conditional input; (ii) an extended configuration, which introduced multi-biomarker conditioning by adding complementary biomarkers (CD16, CD68, and SOX10); and (iii) a full configuration, which incorporates a Cross-Gated attention mechanism alongside the three conditional inputs.

**TABLE II:**
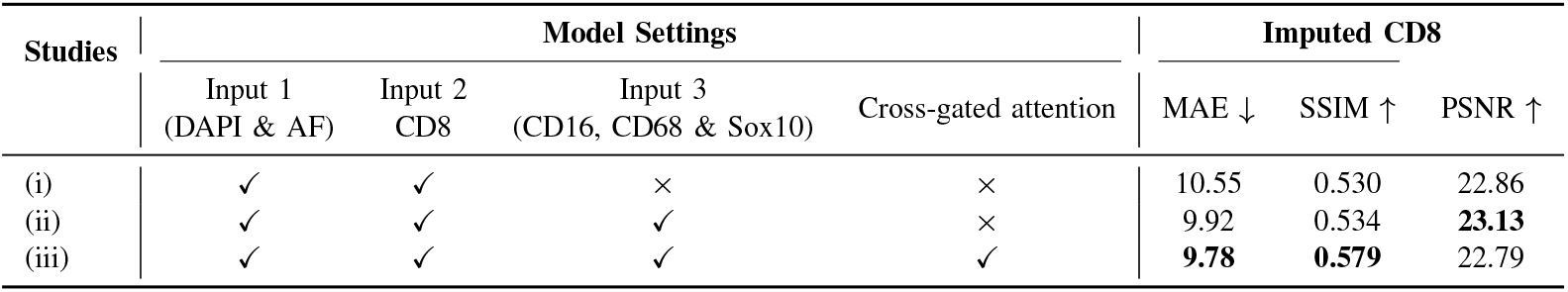
Ablation study.

It was observed that including additional biomarker context (ii) improves SSIM by 0.7%, PSNR by 1.2% and reduces MAE by 6.0%. Furthermore, using the cross-gated attention mechanism significantly improved the performance with a 7.3% reduction in MAE and a 9.2% increase in SSIM compared to the baseline. This suggests that the mechanism effectively enhances feature representation by selectively emphasising informative features while suppressing irrelevant ones and thereby enhances structural reconstruction quality

Furthermore, we note that the inclusion of the Cross-Gated attention mechanism led to a 1.5% drop in PSNR. This may be attributed to the different feature aggregation strategies employed by the two methods. Specifically, in configuration (ii), which excludes cross-gated attention mechanism, the conditional features are directly summed. This simplistic fusion can cause dominant features to suppress more subtle immune-related cues, such as CD8 expression. While this dominance can lead to higher pixel-wise fidelity, it results in poor structural balance, limiting accurate semantic reconstruction. The Cross-Gated attention mechanism alleviates this feature imbalance by adaptively weighting each conditional input based on its contextual relevance, improving structural fidelity.

These findings highlight the importance of incorporating multiple conditional inputs and the Cross-Gated attention mechanism in OS2CR-Diff to effectively integrate these features for improved imputation performance.

### C. CD8^+^T cells segmentation and detection

Detection and segmentation of CD8+T cells play a central role in predicting immunotherapy response, as they serve as key biomarkers of cytotoxic T lymphocyte activity. Therefore, we evaluated the performance of the proposed OS2CR-Diff with OSIM (MAS) for the downstream task of detecting and segmenting CD8+T cells to assess its ability to preserve biologically meaningful features and reduce false-positive rates.

As shown in Table III, the proposed OS2CR-Diff significantly outperforms MAS, which overestimates the true cell count of 20.52 by producing 26.29 cells, whereas OS2CR-Diff generates 17.90 cells, offering a much closer approximation. This implies that OS2CR-Diff more effectively preserves critical cellular features and reduces synthesis artifacts, as illustrated in Fig. 3, resulting in improved accuracy and reliability in CD8+T cells quantification, which is essential for downstream immunological analyses and therapeutic decisionmaking. In precision-critical medical imaging applications, minimising false positives is vital, as overestimation can lead to inaccurate assessments of immune status with potential clinical consequences.

**TABLE III:** Comparison of average segmented CD8^+^T cells counts between OSIM (MAS) and OS2CR-Diff.

| Study | Cell Counts<br>(Real) | Cell Counts<br>(Imputed) | Dice Coefficient ↑ |
| --- | --- | --- | --- |
| MAS | 20.52 | 26.29 (+5.77) | 0.201 |
| **Ours** | 20.52 | **17.90 (−2.62)** | **0.236** |

**Fig. 3:**
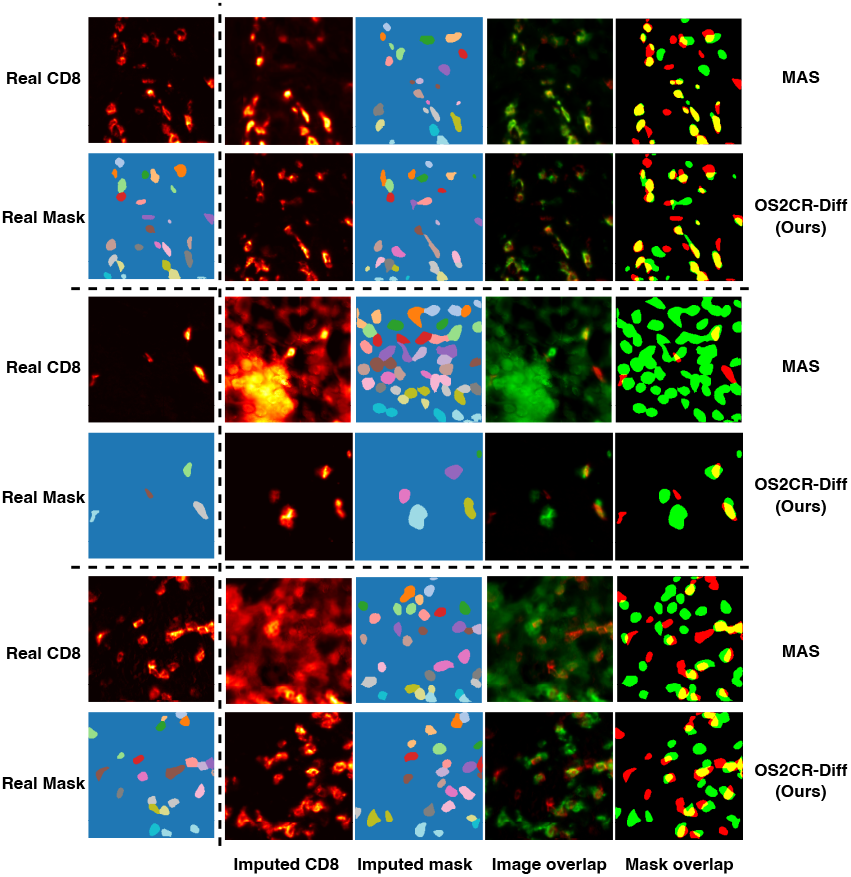
Visualisation of original and imputed images for CD8 segmentation masks and overlap images comparing MAS and OS2CR-Diff.

Moreover, our approach achieves a 6.3% improvement in the Dice coefficient over MAS, indicating enhanced segmentation accuracy. This performance gain underscores the superior reliability of our method in identifying CD8+T cells, an essential cytoplasmic biomarker, thereby strengthening the robustness of our method in clinical diagnostics and immunotherapy evaluation.

## V. Conclusion

In this study, we introduce OS2CR-Diff, a self-refining diffusion framework designed to significantly enhance cytoplasmic biomarker imputation in multiplex immunofluorescence imaging. Our approach integrates a conditional feature fusion block with cross-gated attention mechanisms and velocity prediction parameterisation to iteratively refine outputs from a one-step inference model. The effectiveness of OS2CR-Diff is validated on a melanoma dataset, where it consistently outperforms state-of-the-art methods in CD8 imputation. Notably, it achieves lower false positive rates in CD8^+^T cell counting, thereby providing clinically reliable images for assessing immunotherapy response.

## Notes

### Competing Interest Statement

The authors have declared no competing interest.

## References

[1] Passaro, M. Al Bakir, E. G. Hamilton, M. Diehn, F. André, S. Roy-Chowdhuri, G. Mountzios, I. I. Wistuba, C. Swanton, and S. Peters, “Cancer biomarkers: Emerging trends and clinical implications for personalized treatment,” Cell, vol. 187, no. 7, pp. 1617–1635, 2024.

[2] X. Sun and P. D. Kaufman, “Ki-67: more than a proliferation marker,” Chromosoma, vol. 127, no. 2, pp. 175–186, 2018, Epub 2018 Jan 10.

[3] F. Castro, A. P. Cardoso, R. M. Gonçalves, K. Serre, and M. J. Oliveira, “Interferon-gamma at the crossroads of tumor immune surveillance or evasion,” Frontiers in Immunology, vol. Volume 9 -2018, 2018.

[4] H. Liang, J. Huang, H. Li, W. He, X. Ao, Z. Xie, Y. Chen, Z. Lv, L. Zhang, Y. Zhong, X. Tan, G. Han, J. Zhou, N. Qiu, M. Jiang, H. Xia, Y. Zhan, L. Jiao, J. Ma, D. Radisky, J. Huang, and X. Zhang, “Spatial proximity of CD8(+) T cells to tumor cells predicts neoadjuvant therapy efficacy in breast cancer,” NPJ Breast Cancer, vol. 11, no. 1, pp. 13, 2025, Epub 2025 Feb 10.

[5] M. Sade-Feldman, K. Yizhak, S. L. Bjorgaard, J. P. Ray, C. G. de Boer, R. W. Jenkins, D. J. Lieb, J. H. Chen, D. T. Frederick, M. Barzily-Rokni, S. S. Freeman, A. Reuben, P. J. Hoover, A.-C. Villani, E. Ivanova, A. Portell, P. H. Lizotte, A. R. Aref, J.-P. Eliane, M. R. Hammond, H. Vitzthum, S. M. Blackmon, B. Li, V. Gopalakrishnan, S. M. Reddy, Z. A. Cooper, C. P. Paweletz, D. A. Barbie, A. Stemmer-Rachamimov, K. T. Flaherty, J. A. Wargo, G. M. Boland, R. J. Sullivan, G. Getz, and N. Hacohen, “Defining T cell states associated with response to checkpoint immunotherapy in melanoma,” Cell, vol. 175, no. 4, pp. 998–1013.e20, 2018.

[6] G. Kumar, R. K. Pandurengan, E. R. Parra, K. Kannan, and C. Haymaker, “Spatial modelling of the tumor microenvironment from multiplex immunofluorescence images: methods and applications,” Frontiers in Immunology, vol. 14, pp. 1288802, 2023.

[7] D. Locke and C. C. Hoyt, “Companion diagnostic requirements for spatial biology using multiplex immunofluorescence and multispectral imaging,” Frontiers in Molecular Biosciences, vol. 10, 2023.

[8] C. Hoyt, “Multiplex immunofluorescence and multispectral imaging: Forming the basis of a clinical test platform for immuno-oncology,” Frontiers in Molecular Biosciences, vol. 8, pp. 674747, 2021.

[9] J. Goodfellow, J. Pouget-Abadie, M. Mirza, B. Xu, D. Warde-Farley, S. Ozair, A. C. Courville, and Y. Bengio, “Generative adversarial nets,” in Neural Information Processing Systems, 2014.

[10] A. Dosovitskiy, L. Beyer, A. Kolesnikov, D. Weissenborn, X. Zhai, T. Unterthiner, M. Dehghani, M. Minderer, G. Heigold, S. Gelly, J. Uszkoreit, and N. Houlsby, “An image is worth 16×6 words: Transformers for image recognition at scale,” CoRR, vol. abs/2010.11929, 2020.

[11] X. Li, P. Rana, T. N. Gide, N. A. Adegoke, Y. Mao, G. Long, R. A. Scolyer, S. Berkovsky, E. Coiera, J. S. Wilmott, and S. Liu, “Stain imputation in multiplex immunofluorescence imaging (SIMIF) based on random channel-wise masking,” in 2025 IEEE 22nd International Symposium on Biomedical Imaging (ISBI), 2025, pp. 1–5.

[12] Z. Zhou, Y. Jiang, Z. Sun, T. Zhang, W. Feng, G. Li, R. Li, and L. Xing, “Virtual multiplexed immunofluorescence staining from nonantibody-stained fluorescence imaging for gastric cancer prognosis,” EBioMedicine, vol. 107, pp. 105287, 2024.

[13] J. Kapuscinski, “Dapi: a dna-specific fluorescent probe,” Biotechnic & Histochemistry, vol. 70, no. 5, pp. 220–233, 1995, PMID: 8580206.

[14] S. Srinivasan, C. Zhu, and A. C. McShan, “Structure, function, and immunomodulation of the CD8 co-receptor,” Frontiers in Immunology, vol. Volume 15 - 2024, 2024.

[15] G.-H. Liu, A. Vahdat, D.-A. Huang, E. A. Theodorou, W. Nie, and A. Anandkumar, “I2sb: Image-to-image schrödinger bridge,” in International Conference on Machine Learning, 2023.

[16] J. Ho, A. Jain, and P. Abbeel, “Denoising diffusion probabilistic models,” CoRR, vol. abs/2006.11239, 2020.

[17] Y. Zhou, T. Chen, J. Hou, H. Xie, N. C. Dvornek, S. K. Zhou, D. L. Wilson, J. S. Duncan, C. Liu, and B. Zhou, “Cascaded multi-path shortcut diffusion model for medical image translation,” Medical Image Analysis, vol. 98, pp. 103300, 2024.

[18] O. Ronneberger, P. Fischer, and T. Brox, “U-net: Convolutional networks for biomedical image segmentation,” ArXiv, vol. abs/1505.04597, 2015.

[19] T. Salimans and J. Ho, “Progressive distillation for fast sampling of diffusion models,” CoRR, vol. abs/2202.00512, 2022.

[20] L. Xue, X. Li, and N. L. Zhang, “Not all attention is needed: Gated attention network for sequence data,” CoRR, vol. abs/1912.00349, 2019.

[21] M. Shaban, W. Lassoued, K. Canubas, S. Bailey, Y. Liu, C. Allen, J. Strauss, J. L. Gulley, S. Jiang, F. Mahmood, G. Zaki, and H. A. Sater, “Deep learning model imputes missing stains in multiplex images,” bioRxiv, 2023.

[22] M. M. Zafar, Z. Rauf, A. Sohail, A. R. Khan, M. Obaidullah, S. H. Khan, Y. S. Lee, and A. Khan, “Detection of tumour infiltrating lymphocytes in CD3 and CD8 stained histopathological images using a two-phase deep cnn,” Photodiagnosis and Photodynamic Therapy, vol. 37, pp. 102676, 2022.

[23] Saharia, W. Chan, H. Chang, C. A. Lee, J. Ho, T. Salimans, D. J. Fleet, and M. Norouzi, “Palette: Image-to-image diffusion models,” CoRR, vol. abs/2111.05826, 2021.

[24] Z. Yaseen, T.N. Gide, J.W. Conway, A.J. Potter, C. Quek, A.M. Hong, G.V. Long, R.A. Scolyer, and J.S. Wilmott, “Validation of an accurate automated multiplex immunofluorescence method for immuno-profiling melanoma,” Frontiers in Molecular Bioscience, vol. 9, no. e810858, 2022.

[25] T. N. Gide, C. Quek, A. M. Menzies, A. T. Tasker, P. Shang, J. Holst, J. Madore, S. Y. Lim, R. Velickovic, M. Wongchenko, Y. Yan, S. Lo, M. S. Carlino, A. Guminski, R. P. M. Saw, A. Pang, H. M. McGuire, U. Palendria, J. F. Thompson, H. Rizos, I. P. Da Silva, M. Batte, A. Scolyer, G. V. Long, and J. S. Wilmott, “Distinct immune cell populations define response to anti-PD-1 monotherapy and anti-PD-1/anti-ctla-4 combined therapy,” Cancer Cell, vol. 35, no. 2, pp. 238–255, 2019.

[26] N. Otsu, “A threshold selection method from gray-level histograms,” IEEE Transactions on Systems, Man, and Cybernetics, vol. 9, no. 1, pp. 62–66, 1979.

[27] Y. Li, S. Yang, X. Wu, S. He, and S. K. Zhou, “Taming stable diffusion for mri cross-modality translation,” in 2024 IEEE International Conference on Bioinformatics and Biomedicine (BIBM), 2024, pp. 2134– 2141.

[28] Saharia, W. Chan, S. Saxena, L. Li, J. Whang, E. L. Denton, K. S. Ghasemipour, B. K. Ayan, S. S. Mahdavi, R. G. Lopes, Salimans, J. Ho, D. J. Fleet, and M. Norouzi, “Photorealistic text-to-image diffusion models with deep language understanding,” ArXiv, vol. abs/2205.11487, 2022.

[29] C. J. Willmott and K. Matsuura, “Advantages of the mean absolute error (MAE) over the root mean square error (RMSE) in assessing average model performance,” Climate Research, vol. 30, pp. 79–82, 2005.

[30] A. Horé and D. Ziou, “Image quality metrics: Psnr vs. ssim,” in 2010 20th International Conference on Pattern Recognition, 2010, pp. 2366– 2369.

[31] M. Pachitariu, M. Rariden, and C. Stringer, “Cellpose-sam: superhuman generalization for cellular segmentation,” bioRxiv, 2025.

[32] L. R. Dice, “Measures of the amount of ecologic association between species,” Ecology, vol. 26, no. 3, pp. 297–302, 1945.

